# The *Drosophila* metallopeptidase *superdeath* decouples apoptosis from the activation of the ER stress response

**DOI:** 10.1101/620492

**Authors:** Rebecca A.S. Palu, Clement Y. Chow

**Author notes:** Corresponding author: Clement Y. Chow, 15 N 2030 E, Rm 5200, Eccles Institute of Human Genetics Salt Lake City, UT 84112.

## Abstract

Endoplasmic reticulum (ER) stress-induced apoptosis is a primary cause and modifier of degeneration in a number of genetic disorders. Understanding how genetic variation between individuals influences the ER stress response and subsequent activation of apoptosis could improve individualized therapies and predictions of outcomes for patients. In this study, we find that the uncharacterized, membrane-bound metallopeptidase *CG14516* in *Drosophila melanogaster*, which we rename as *<u>SUP</u>pressor of <u>ER</u> stress-induced <u>DEATH</u>* (*superdeath*), plays a role in modifying ER stress-induced apoptosis. We demonstrate that loss of *superdeath* reduces apoptosis and degeneration in the *Rh1*^*G69D*^ model of ER stress through the JNK signaling cascade. This effect on apoptosis occurs without altering the activation of the unfolded protein response (IRE1 and PERK), suggesting that the beneficial pro-survival effects of this response are intact. Furthermore, we show that *superdeath* functions epistatically upstream of *CDK5*, a known JNK-activated pro-apoptotic factor in this model of ER stress. We demonstrate that *superdeath* is not only a modifier of this particular model, but functions as a general modifier of ER stress-induced apoptosis across different tissues and ER stresses. Finally, we present evidence of Superdeath localization to the endoplasmic reticulum membrane. While similar in sequence to a number of human metallopeptidases found in the plasma membrane and ER membrane, its localization suggests that *superdeath* is orthologous to *ERAP1/2* in humans. Together, this study provides evidence that *superdeath* is a link between stress in the ER and activation of cytosolic apoptotic pathways.

**SIGNIFICANCE STATEMENT:** Genetic diseases display a great deal of variability in presentation, progression, and overall outcomes. Much of this variability is caused by differences in genetic background among patients. One process that commonly modifies degenerative disease is the endoplasmic reticulum (ER) stress response. Understanding the genetic sources of variation in the ER stress response could improve individual diagnosis and treatment decisions. In this study, we characterized one such modifier in *Drosophila melanogaster*, the membrane-bound metallopeptidase *CG14516* (*superdeath*). Loss of this enzyme suppresses a model of ER stress-induced degeneration by reducing cell death without altering the beneficial activation of the unfolded protein response. Our findings make *superdeath* and its orthologues attractive therapeutic targets in degenerative disease.

## INTRODUCTION

Endoplasmic reticulum (ER) stress-induced apoptosis is a primary or contributing cause of degeneration in a wide variety of diseases such as type 2 diabetes and Alzheimer’s Disease (1–3). Reducing stress-induced apoptosis could be the key to slowing the progression of these diseases, and indeed a major focus of therapeutic development is to identify compounds that can inhibit apoptosis. Therapeutics that target cell death without impacting the beneficial survival pathways activated by the ER stress response are essential in the treatment of degenerative diseases.

ER stress occurs when protein folding is disrupted, leading to the accumulation of misfolded proteins in the ER. ER stress activates the Unfolded Protein Response (UPR), a massive transcriptional response that, if successful, will return the ER and cell to homeostasis (4). Under conditions of chronic or extreme stress, the UPR may eventually induce apoptosis (5). In *Drosophila*, this is primarily through the activation of the Jun Kinase (JNK) signaling cascade downstream of the cell cycle regulator CDK5 (5, 6). The mechanism through which CDK5 is activated by ER stress and the UPR is unknown.

The ER stress response is strongly influenced by genetic variation (7–10). In a previous study, we modeled the impact of genetic variation on the ER stress response by overexpressing mutant rhodopsin (*Rh1*^*G69D*^) in the developing *Drosophila* eye (9). We crossed this model into the ~200 genetic backgrounds of the *Drosophila* Genetic Reference Panel (DGRP) (9, 11). We measured retinal degeneration and performed a genome-wide association analysis to identify modifier variation that is associated with differences in degeneration. We generated a list of 84 conserved candidate modifier genes, approximately 50% of which have known roles in apoptotic pathways and/or the ER stress response (9). By characterizing these modifiers, we can learn more about the pathogenesis and progression of ER stress-related diseases.

Here, we report a novel function for one of these modifiers, the *Drosophila* metallopeptidase *CG14516*, which we rename *<u>SUP</u>pressor of <u>ER</u> stress*-*induced <u>DEATH</u>* (*superdeath)* (12). We demonstrate that, in this *Rh1*^*G69D*^ model of ER stress, loss of *superdeath* results in partial rescue of degeneration. This reduced degeneration is accompanied by reduced apoptosis and JNK signaling, but occurs in the absence of any detectable changes in activation of the ER stress sensors IRE1 or PERK, suggesting that *superdeath* lies downstream of the UPR in the activation of apoptosis. Epistasis experiments indicate that *superdeath* lies genetically upstream of *CDK5* and that the changes observed in degeneration are due to reduced activation of CDK5. While this gene is orthologous to a number of mammalian metallopeptidases, we show that superdeath is localized to the ER, suggesting that it is functionally related to the ER-associated proteases 1 and 2 (ERAP1 and ERAP2) (13). Our results suggest that inhibition of Superdeath/ERAP1/ERAP2 would reduce apoptosis levels under conditions of ER stress, while retaining the beneficial effects of UPR activation, making it a valuable target for therapeutic development.

## RESULTS

### A SNP in *CG14516* is associated with variation in expression and degeneration in the *Rh1*^*G69D*^ model of ER stress

In a previous study, we examined the impact of genetic variation on ER stress-induced apoptosis using a model of ER stress in which we over-expressed mutant, misfolding rhodopsin in the developing eye imaginal disc using the *GAL4*/*UAS* system (*GMR*-*GAL4*>*UAS*-*Rh1*^*G69D*^) (9, 14). We crossed this model of retinal degeneration and ER stress into the DGRP, a collection of ~200 wild-derived isogenic strains (11). We performed a genome-wide associated analysis to identify candidate modifier genes of ER stress and degeneration. One of these candidate genes is *CG14516*, a previously uncharacterized membrane-bound metallopeptidase. A SNP in the first intron of *CG14516* is significantly associated with eye size in the *Rh1*^*G69D*^ model (P = 2.24×10^−5^, Fig 1A). Strains carrying the major “A” allele have larger, less degenerate eyes (21922 ± 2279 pixels) compared to those carrying the minor “T” allele (19863 ± 2873 pixels). The minor “T” allele is also associated with a small, but significant increase in expression of *CG14516* in adult females (9.09 ± 0.24 log_2_ [FPKM +1]) compared to the major “A” allele (8.84 ± 0.30 log_2_ [FPKM +1], P = 4.92×10^−3^, Fig 1B) (15). Importantly, overall expression of *CG14516* in adult females is inversely correlated with eye size in the presence of *Rh1*^*G69D*^ (r = −0.25, P = 0.0013, Fig 1C, D). These data show that reduced expression of *CG14516* is associated with a decrease in *Rh1*^*G69D*^-induced degeneration, suggesting that loss of *CG14516* function should reduce ER stress-induced degeneration. We have therefore named this gene *<u>SUP</u>pressor of <u>ER</u> stress*-*induced <u>DEATH</u>* (*superdeath*).

**Figure 1.**
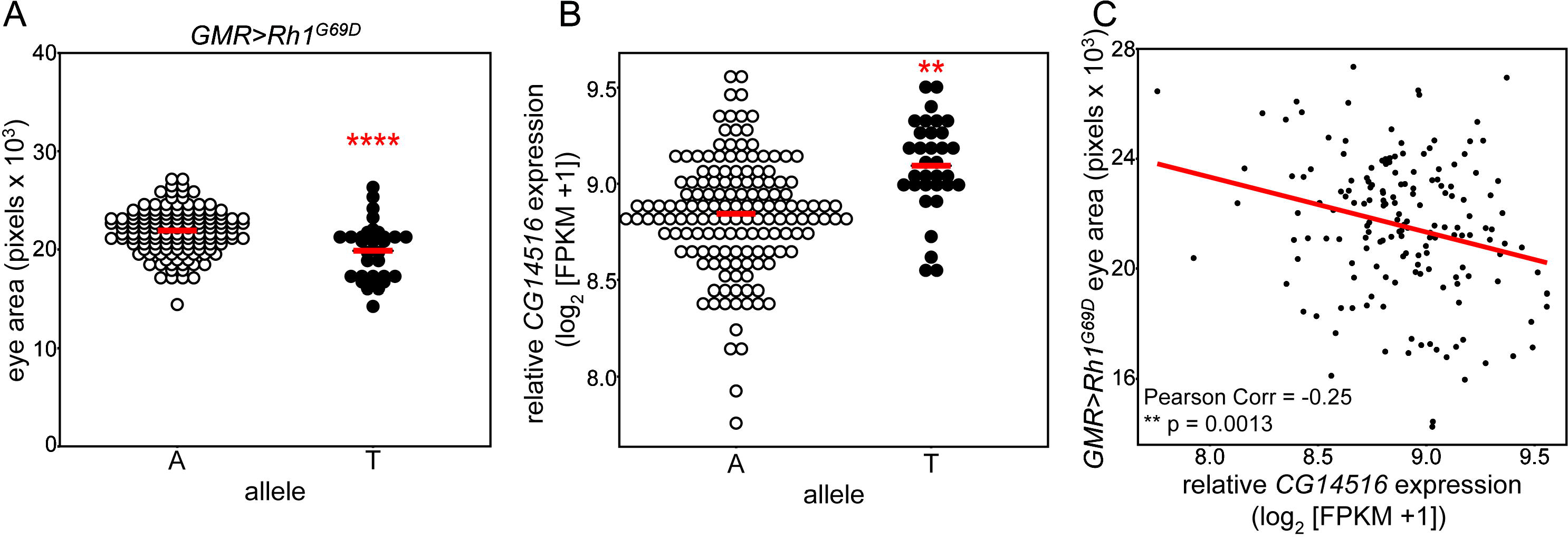
A SNP in *CG14516* is associated with changes in expression and degeneration. Variation in sequence and expression of the *Drosophila melanogaster* gene *CG14516* is associated with eye size in the *Rh1*^*G69D*^ model of ER stress. **A.** The 3R:24966022 SNP in *CG14516* (BDGP R5/dm3) is associated with *Rh1*^*G69D*^-induced degeneration. *Rh1*^*G69D*^ DGRP eye size is plotted by allele identity. Strains carrying the minor “T” allele (19863 ± 2873 pixels, N = 32) have significantly smaller eyes than those carrying the major “A” allele (21922 ± 2279 pixels, N = 136). **B.** Expression of *CG14516* in strains carrying either the “A” or the “T” allele was determined from available RNA sequencing data in adult females (15). *CG14516* levels was significantly increased in strains carrying the minor “T” allele (9.09 ± 0.24 units, N = 33) as compared to those carrying the major “C” allele (8.84 ± 0.30 units, N = 147). **C.** Eye size in the *Rh1*^*G69D*^ DGRP strains is inversely correlated with *CG14516* expression levels in adult females (r = −0.25, N = 167, P = 0.0013). Raw data for **A** and **C** were taken from Chow *et.al* 2016 and Huang *et.al* 2015, respectively. Values are average ± SD. ** P < 0.005, **** P <0.00005.

### Loss of *superdeath* expression rescues *Rh1*^*G69D*^-induced apoptosis and degeneration

To test the impact of *superdeath* expression on *Rh1*^*G69D*^-induced degeneration, we expressed an RNAi construct targeting *superdeath* in the presence of the *Rh1*^*G69D*^ model of ER stress (*Rh1*^*G69D*^/*superdeathi*). Ubiquitous expression of this RNAi construct results in a 75% reduction in *superdeath* expression (0.278 ± 0.072 relative to controls at 1.00 ± 0.16, P = 1.77×10^−4^, Fig S1A). We found that in the absence of *superdeath*, eye size is significantly increased (15299 ± 1658 pixels) compared to a control that is only expressing *Rh1*^*G69D*^ (*Rh1*^*G69D*^/control) (11942 ± 473 pixels, P = 8.15×10^−6^, Fig 2A). We observe a slight increase in eye size when we reduce expression of *superdeath* in wild-type eyes compared to controls, but no qualitative difference (28867 ± 1566 pixels vs. 25968 ± 1026 pixels in controls, P = 1.16^−4^, Fig 2B).

**Figure 2.**
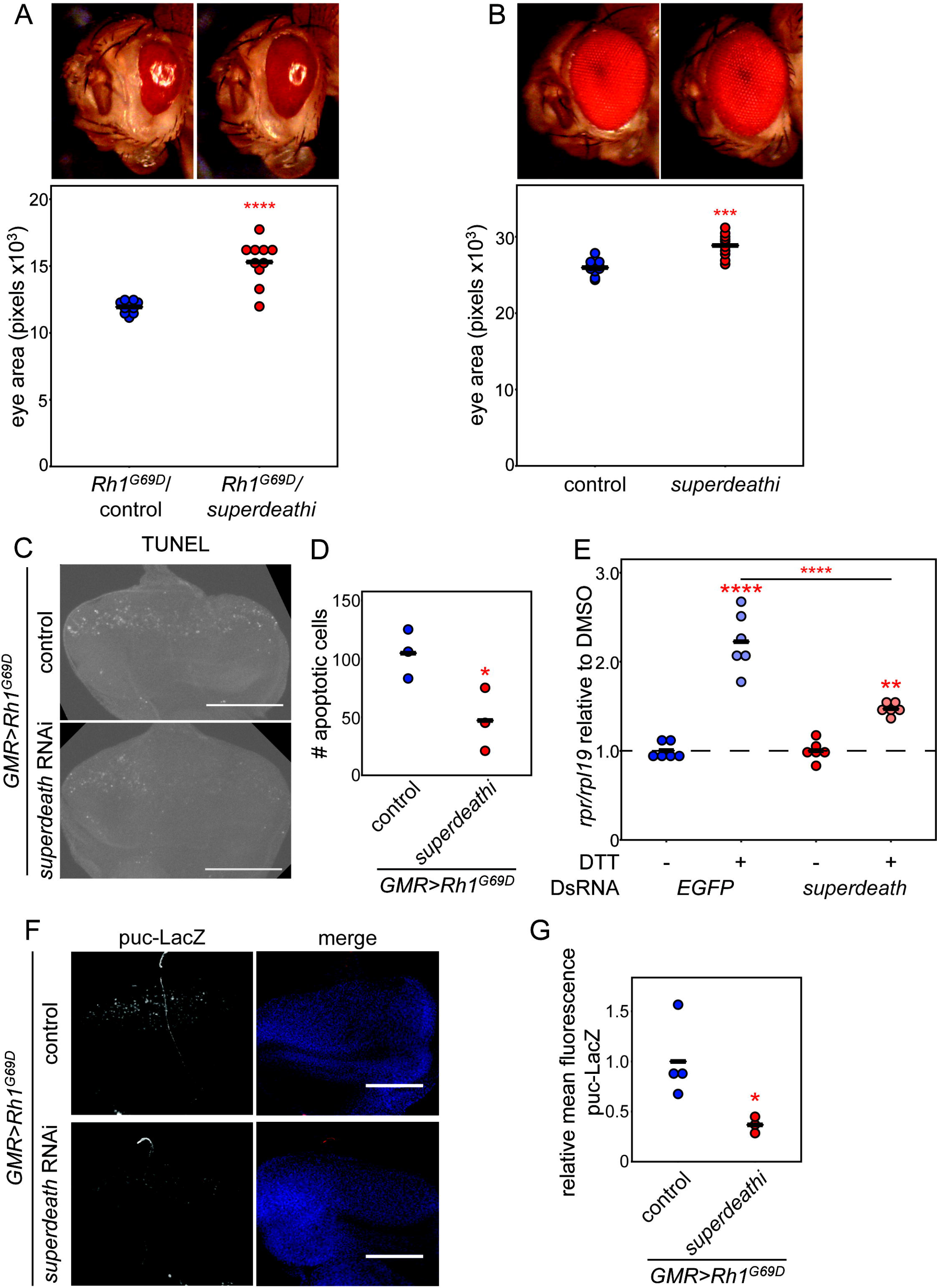
Loss of *superdeath* reduces ER stress-associated apoptosis. Reducing expression of *superdeath* reduces apoptosis and degeneration in models of ER stress. **A.** Degeneration caused by overexpression of *Rh1*^*G69D*^ is partially rescued by RNAi-mediated knockdown of *superdeath* expression (15299 ± 1658 pixels, N = 10 in *Rh1*^*G69D*^/*superdeathi* flies as compared to 11942 ± 473 pixels, N = 10 in *Rh1*^*G69D*^/controls). **B.** Eye size also showed a small increase when the *superdeath* RNAi construct was expressed in a wild-type background (28867 ± 1566 pixels, N = 10) as compared to controls (25968 ± 1026 pixels, N = 10), but no qualitative differences were observed. **C.** *Rh1*^*G69D*^/*superdeathi* eye imaginal discs display reduced apoptosis compared to *Rh1*^*G69D*^/controls as measured by TUNEL staining. **D.** S2 cells treated with DsRNA against *EGFP* increased expression of the apoptotic gene *rpr* after 7.5 hours of DTT exposure (2.23 ± 0.33, N = 6) as compared to DMSO-treated control cells (1.00 ± 0.09, N = 6). Activation of *rpr* expression was significantly reduced in S2 cells that were treated with DsRNA against *superdeath* (1.47 ± 0.07, N = 6 in DTT-treated cells compared to 1.00 ± 0.11, N = 6 with DMSO). **E.** Activation of JNK signaling was reduced in *Rh1*^*G69D*^/*superdeathi* eye imaginal discs compared to *Rh1*^*G69D*^/controls as determined by expression of *puc*-*LacZ*. **F.** When quantified, LacZ levels were significantly lower in *Rh1*^*G69D*^/*superdeathi* eye discs (0.367 ± 0.082, N = 3) as compared to *Rh1*^*G69D*^/controls (1.00 ± 0.39, N = 4). Values are average ± SD. Scale bars = 0.1 mm. * P < 0.05, ** P < 0.005, **** P < 0.00005.

Because reduced degeneration in the *Rh1*^*G69D*^ model of ER stress is often accompanied by reduced apoptosis (6, 16), we measured cell death in the eye imaginal disc, a developmental structure that will eventually become the adult eye. This tissue is also the site of mutant rhodopsin overexpression and where ER stress and apoptosis are being induced. TUNEL staining indicates that there is reduced apoptosis in the absence of *superdeath* (47 ± 27 cells) compared to *Rh1*^*G69D*^/controls (104 ± 21 cells, P = 0.044, Fig 2C,D). Our findings demonstrate that *superdeath* is required for high levels of ER stress-induced apoptosis and subsequent degeneration. The rescue effect observed upon loss of *superdeath* is likely due to this reduced apoptosis.

To confirm these findings in an alternate system, we treated *Drosophila* S2 cells with DsRNA targeting either *superdeath* or *EGFP* as a control. This resulted in 90% reduction in *superdeath* expression (0.102 ± 0.038 relative to controls at 1.00 ± 0.29, P = 1.9×10^−5^, Fig S1B). S2 cells were then treated with Dithiothreitol (DTT) to induce ER stress. DTT disrupts disulfide bond formation and results in massive protein misfolding, ER stress, and the induction of the UPR (17). We monitored expression of the apoptosis-associated transcript *rpr* to determine if there is a difference in the induction of apoptosis when *superdeath* expression is reduced in S2 cells. rpr is an inhibitor protein that targets the Inhibitor of Apoptosis Proteins (IAPs), enabling caspase activation and the initiation of apoptosis (18, 19). Its expression is upregulated under most apoptosis-inducing conditions (20, 21). As expected, treatment of control S2 cells with DTT results in a ~2.25-fold increase in *rpr* expression (2.23 ± 0.33) compared to cells treated with DMSO (1.00 ± 0.09, P < 10^−7^), indicating that a larger percentage of the DTT-treated cells are undergoing cell death (Fig 2E). In contrast, S2 cells lacking expression of *superdeath* that are treated with DTT display an approximately 1.5-fold increase in *rpr* expression (1.47 ± 0.07) as compared to DMSO-treated cells (1.00 ± 0.11, P = 1.2×10^−3^, Fig 2E). This response is significantly weaker than that seen in control cells (P = 3.5×10^−6^). These results support a role for *superdeath* in apoptosis activation.

### *superdeath* regulates JNK signaling independently of UPR activation

In *Drosophila* models of ER stress, and specifically in this model, apoptosis is initiated through activation of the JNK signaling cascade (6, 22). To determine if JNK signaling is disrupted upon loss of Superdeath activity, we monitored the expression of a known Jun target gene, *puckered* (*puc*). We used an allele of *puc* wherein the coding sequence has been replaced by the coding sequence for *LacZ*, such that *LacZ* expression is driven by the promotor and regulatory sequences that normally govern *puc* expression (23). LacZ levels serve as a direct readout for binding of the *puc* promotor by the Jun transcription factor. As expected, we detected high expression of LacZ in the *Rh1*^*G69D*^/control eye imaginal discs (1.00 ± 0.39, Fig 2F). This expression is significantly reduced in the absence of *superdeath* (0.367 ± 0.082 relative to controls, P = 0.043, Fig. 2F,G). Our findings support a model of reduced signaling through the JNK cascade, which ultimately results in reduced apoptosis.

### Loss of *superdeath* does not impact the activation of UPR signaling pathways

We hypothesized that the reduction in apoptosis and JNK signaling observed in the absence of *superdeath* might be caused by reduced activation of the UPR. We therefore monitored the activation of two of the UPR sensors: IRE1 and PERK. IRE1 is the most conserved of the ER stress sensors. When activated by the accumulation of misfolded proteins, the RNAse domain of IRE1 is responsible for the non-canonical splicing of the mRNA for the transcription factor *Xbp1* (5, 24). The spliced isoform of *Xbp1* is then translated and travels to the nucleus, where it activates the expression of UPR target genes. We monitored IRE1 activity in the eye imaginal disc using an *Xbp1* transgene where the 3’ end of the transcript has been replaced with the coding sequence for *EGFP*, such that EGFP is only expressed under conditions that induce IRE1 activity and *Xbp1* splicing (14, 25–27). IRE1 activity was measured by staining for EGFP in eye imaginal discs dissected from *Rh1*^*G69D*^/control and *Rh1*^*G69D*^/*superdeathi* flies. We also monitored Rh1^G69D^ levels using an antibody against rhodopsin to determine if there are differences in the amount of misfolded protein being expressed. We detected no significant differences in either EGFP (0.905 ± 0.057 relative to 1.00 ± 0.25 in controls) or rhodopsin levels (0.899 ± 0.071 relative to 1.00 ± 0.10 in controls, Fig 3A-C). Similarly, there was no significant difference in *Xbp1* splicing after exposure to DTT in S2 cells treated with DsRNA targeting *EGFP* or *superdeath* (Fig 3D). These results collectively demonstrate that loss of *superdeath* does not influence IRE1 activation in response to ER stress.

**Figure 3.**
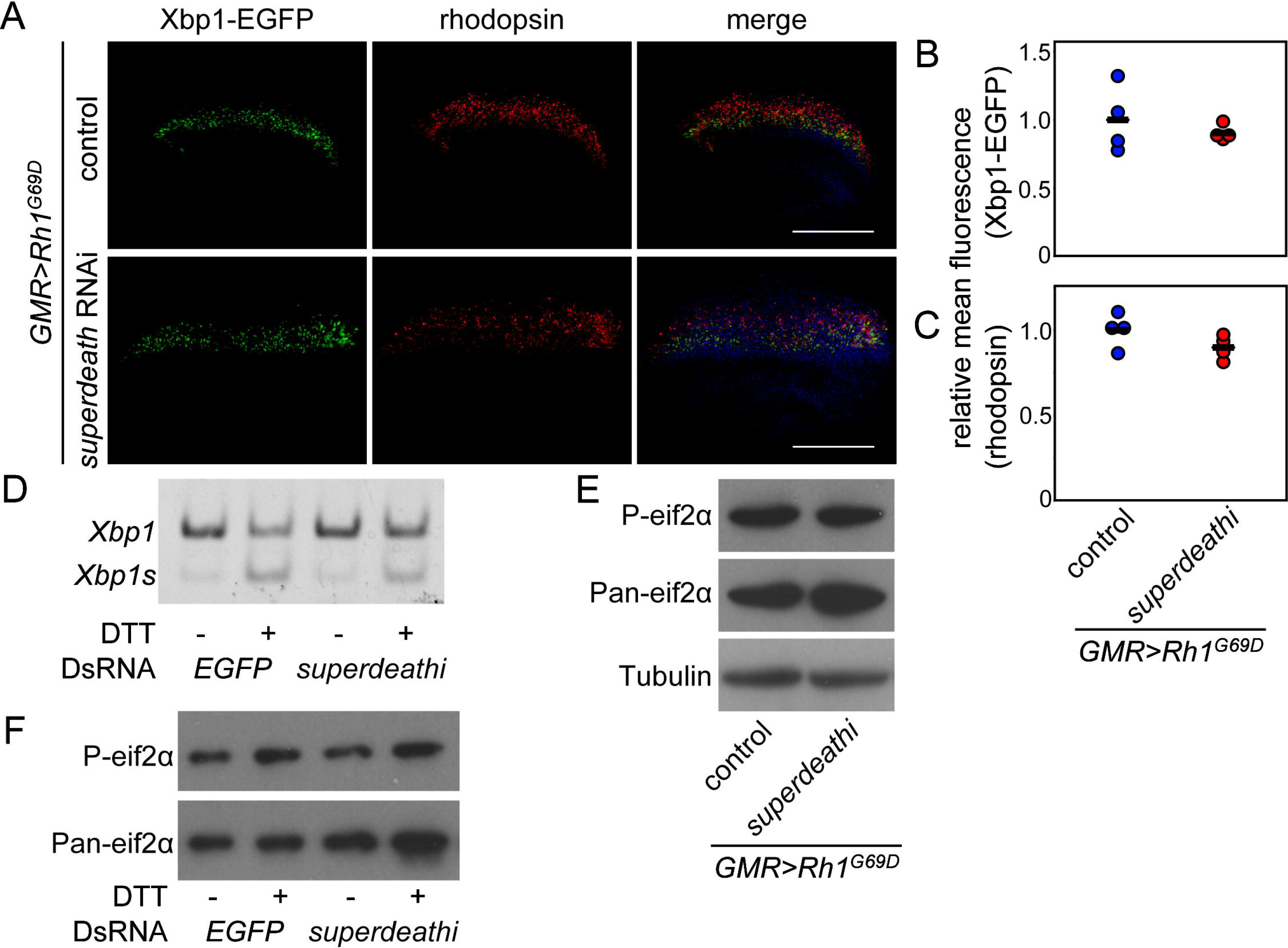
Loss of *superdeath* does not alter IRE1 or PERK activation. Activation of the UPR is not altered by loss of *superdeath* in models of ER stress. **A.** *Rh1*^*G69D*^/*superdeathi* eye discs do not display altered expression of Xbp1-EGFP or rhodopsin as compared to *Rh1*^*G69D*^/controls. Eye discs were dissected from wandering L3 larvae expressing *Rh1*^*G69D*^ and *UAS*-*Xbp1*-*EGFP*, stained for rhodopsin and GFP as an indicator of ER stress, and counterstained with 4’,6-diamidino-2-pheneylindole (DAPI). **B.** Loss of *superdeath* does not significantly alter Xbp1-EGFP expression (0.905 ± 0.057, N = 4) compared to *Rh1*^*G69D*^/controls (1.00 ± 0.25, N = 4). **C.** Rhodopsin levels were also not significantly altered (0.899 ± 0.071, N = 4 relative to 1.00 ± 0.10, N = 4 in controls). **D.** DTT treatment increased *Xbp1* splicing in S2 cells compared to control cells treated with DMSO. This increase was similar in S2 cells treated with DsRNA against either *EGFP* or *superdeath*. **E.** *Rh1*^*G69D*^/*superdeathi* eye discs had similar levels of P-eif2α as compared to *Rh1*^*G69D*^/controls. **F.** DTT treatment in S2 cells increased levels of P-eif2α compared to the control DMSO treatment. This increase was similar to cells treated with DsRNA against either *EGFP* or *superdeath*. Values are average ± SD. Scale bars = 0.1 mm.

The second major sensor of the UPR is the kinase PERK, which, upon activation by the accumulation of misfolded proteins, phosphorylates the translation initiation factor eif2a. This modification reduces the efficiency of canonical translation initiation while allowing for increased translation of select UPR regulators such as the transcription factor ATF4 (5). To assess PERK activity, we monitored eif2a phosphorylation by Western blot of samples isolated from brain-imaginal disc complexes of *Rh1*^*G69D*^/*superdeathi* and *Rh1*^*G69D*^/control larvae. We detected no significant differences in P-eif2a accumulation in these samples, suggesting that PERK activity is also unaffected by reduced expression of *superdeath* (Fig 3E). Phosphorylation of eif2a after DTT treatment is also similar between cells treated with DsRNA against either *EGFP* or *superdeath* (Fig 3F).

Collectively, these results indicate that loss of *superdeath* activity does not impact the UPR and the reduced apoptosis and JNK signaling observed in the absence of *superdeath* is independent of UPR activation or the accumulation of misfolded proteins.

### *superdeath* functions upstream of *CDK5* in ER stress-induced apoptosis

*superdeath* could be regulating general apoptosis signaling or could act more specifically on pathways activated by the UPR. To test if *superdeath* is generally involved in the regulation of cell death, we expressed RNAi targeting *superdeath* in the developing eye imaginal discs of flies overexpressing the cell death initiators *p53* and *rpr*. p53 is primarily activated by the DNA damage response and can initiate apoptosis by transcriptionally activating the IAP inhibitors *rpr*, *grim*, and *hid* (21). *rpr* is activated transcriptionally by p53 and the JNK signaling cascade (20, 21, 23). Overexpression of either of these factors in the eye imaginal disc is sufficient to induce extensive apoptosis and a retinal degenerative phenotype in adult flies (18, 28). We can therefore test the impact of *superdeath* expression on general apoptotic pathways by expressing the RNAi construct targeting *superdeath* in models of *p53* or *rpr* overexpression and evaluating changes in eye degeneration.

We first tested loss of *superdeath* in a model of *p53* overexpression (*p53*/*superdeathi*) to determine whether general cell death pathways are impacted. We found no difference in eye size between *p53*/control (14852 ± 1126 pixels) and *p53*/*superdeathi* flies (15315 ± 1000 pixels, Fig 4A). We next tested *superdeath* function in a model of *rpr* overexpression (*rpr*/*superdeathi*) to see if the function of this gene lies upstream or downstream of the transcriptional program that commonly activates apoptosis. As with *p53*, there was no difference in eye size between *rpr*/control (18953 ± 834 pixels) and *rpr*/*superdeathi* flies (19288 ± 664 pixels, Fig 4B). Our findings suggest that *superdeath* functions upstream of the transcriptional program that initiates apoptosis and in a pathway that is specific to ER stress.

**Figure 4.**
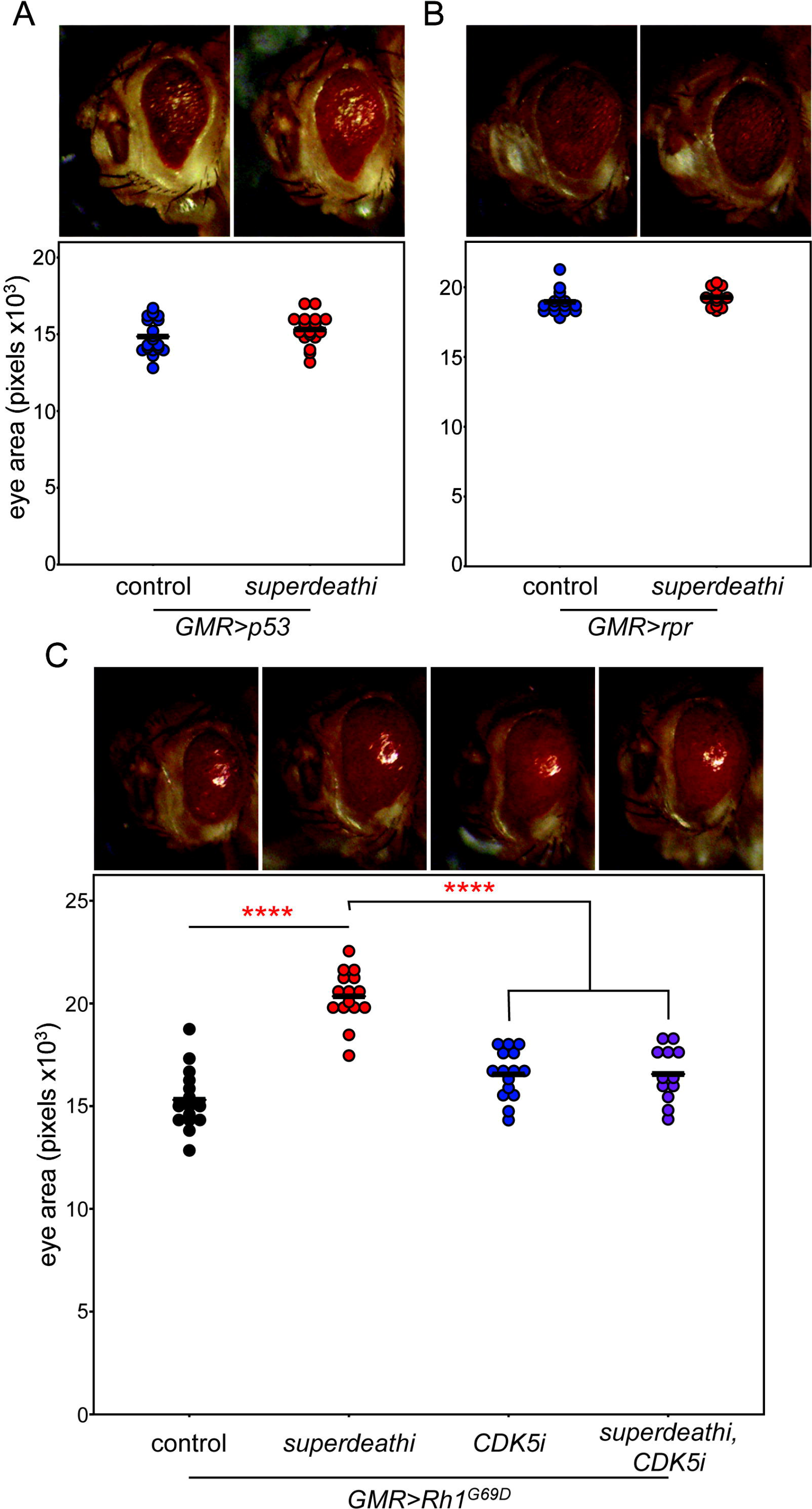
*superdeath* is upstream of *CDK5* in the initiation of ER stress-induced apoptosis. *superdeath* acts specifically in ER stress-associated cell death, upstream of the transcription of apoptotic activators. **A.** Degeneration caused by overexpression of *p53* is not altered by RNAi-mediated knockdown of *superdeath* expression (15315 ± 1000 pixels, N = 18 in *p53*/*superdeathi* flies compared to 14852 ± 1126 pixels, N = 18 in *p53*/controls). **B.** Degeneration caused by overexpression of *rpr* is also unaffected by loss of *superdeath* expression (19288 ± 664 pixels, N = 13 in *rpr/superdeathi* flies compared to 18953 ± 834 pixels, N = 15 in *rpr*/controls). **C.** Degeneration caused by overexpression of Rh1^G69D^ in the absence of both *CDK5* and *superdeath* (16560 ± 1320 pixels, N = 12) does not significantly differ from degeneration in the absence of *CDK5* alone (16552 ± 1179 pixels, N = 15). Degeneration is qualitatively improved by RNAi-mediated knockdown of *CDK5* expression although eye size is not significantly increased compared to Rh1^G69D^/controls (15307 ± 1482 pixels, N = 15). This is in contrast to the significant increase in eye size when *superdeath* expression is reduced (20346 ± 1292 pixels, N = 15). Values are average ± SD. **** P < 0.00005.

Activation of JNK-induced apoptosis in the *Rh1*^*G69D*^ model of ER stress is regulated by the cyclin-dependent kinase CDK5. Loss of *CDK5* leads to reduced activation of apoptosis without altering the activation of the UPR (6). These molecular changes are accompanied by qualitative improvements in eye size, morphology, and pigmentation (6, 9). We hypothesized that *superdeath* might function in this pathway. To test this, we expressed RNAi constructs targeting *CDK5* and *superdeath*, individually and concurrently, in the developing eye imaginal discs expressing the misfolded Rh1^G69D^ protein. We monitored degeneration using eye size in adult flies. As expected, loss of *superdeath* (*Rh1*^*G69D*^/*superdeathi*) results in a substantial and significant increase in eye size as compared to the *Rh1*^*G69D*^/controls (20345 ± 1292 pixels vs. 15307 ± 1482 pixels in controls, P <0.00005, Fig 4C). In line with previous reports (6, 9), expressing the misfolded Rh1^G69D^ protein and RNAi against *CDK5* (*Rh1*^*G69D*^/*CDK5i*) results in a qualitative rescue and improvement in eye morphology, but no change in eye size as compared to controls (16552 ± 1179 pixels, P = 0.060, Fig 4C). Because the phenotype associated with loss of *CDK5* is distinguishable from the phenotype associated with loss of *superdeath*, we can perform an epistasis experiment to determine which of these genes lies downstream of the other. We found that flies simultaneously expressing both RNAi against *CDK5* and *superdeath* (*Rh1*^*G69D*^/*CDK5i*-*superdeathi*) display similar phenotypes to *Rh1*^*G69D*^/*CDK5i* flies, with no quantitative improvement in eye size as compared to controls (16560 ± 1320 pixels, P = 0.081, Fig 4C). We concluded that *superdeath* must operate upstream of *CDK5* to regulate the activation of JNK signaling and apoptosis.

### *superdeath* regulates ER stress-induced apoptosis in multiple tissues

Because CDK5 and JNK signaling occur across different ER stress conditions, we tested whether *superdeath* can act as a modifier across different tissues and methods of ER stress initiation. The wing imaginal disc is a developmental structure that will eventually mature and become the adult wing. Wing imaginal disc expression of a misfolded protein such as Rh1^G69D^ using the *MS1096*-*GAL4* driver (wing-*Rh1*^*G69D*^/control) induces ER stress and apoptosis, resulting in a small, degenerate wing that fails to unfold upon eclosion (Fig 5A). Concurrent expression of *Rh1*^*G69D*^ and RNAi against *superdeath* (wing-*Rh1*^*G69D*^/*superdeathi*) results in partial rescue of the degenerate wing phenotype (Fig 5A), similar to what was observed for the eye. This is also reflected in wing area, which increases from wing-*Rh1*^*G69D*^/*superdeathi* (94627 ± 26363 pixels) flies compared to wing-*Rh1*^*G69D*^/controls (73931 ± 19988 pixels, P = 0.044, Fig 5A) flies.

**Figure 5.**
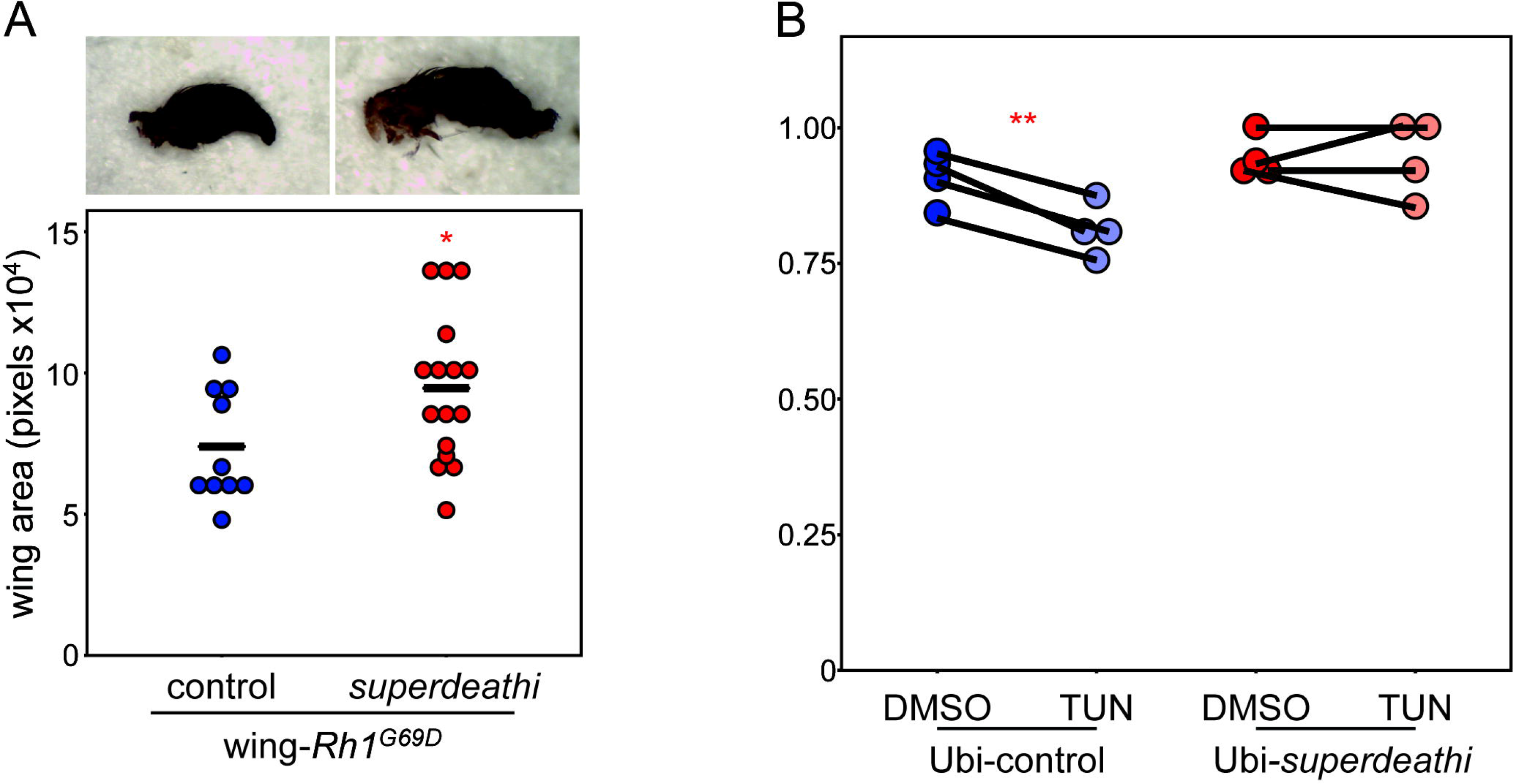
*superdeath* is a general regulator of ER stress-induced cell death. The ER stress response and subsequent apoptosis is subject to regulation by *superdeath* across ER stress conditions. **A.** Loss of *superdeath* partially rescues the vestigial wing phenotype caused by expression of *Rh1*^*G69D*^ in the wing disc (94627 ± 26363 pixels, N = 16 in wing-*Rh1*^*G69D*^/*superdeathi* vs. 73931 ± 19988 pixels, N = 10 in wing-*Rh1*^*G69D*^/controls). **B.** Larvae with ubiquitous knockdown of *superdeath* are significantly more resistant to tunicamycin-induced ER stress than control larvae. Four paired experimental replicates are shown, representing a combined total of N = 113 DMSO-treated and N = 130 TUN-treated Ubi-control larvae, and N = 112 DMSO-treated and N = 127 TUN-treated Ubi-*superdeathi* larvae. Values are average ± SD. * P < 0.05, ** P < 0.005.

To determine whether *superdeath* also responds to diverse mechanisms of initiating ER stress, we ubiquitously expressed RNAi targeting *superdeath* using the *Tub*-*GAL4* driver (Ubi-*superdeathi*). We then exposed Ubi-*superdeathi* larvae along with controls expressing only *Tub*-*GAL4* (Ubi-control) to tunicamycin or DMSO for four hours. Tunicamycin inhibits N-linked glycosylation in the ER, inducing a massive ER stress response through the UPR (29). This five-hour treatment is sufficient to significantly activate the UPR in the larvae, leading to developmental delay and lethality (15, 30). We found that Ubi-*superdeathi* larvae are significantly more resistant to tunicamycin-induced lethality as compared to Ubi-control larvae (Fig 5B, Table S1). We concluded from these results, and the S2 cell data above, that *superdeath* is generally important for the regulation of degeneration and apoptosis downstream of the initiation of ER stress.

### Superdeath is localized the endoplasmic reticulum

Based on sequence analysis, Superdeath is hypothesized to contain a single transmembrane domain near the N-terminal end of the protein, with the active site localized to the cytoplasm of the cell (31). Based on our results above, we hypothesized that it could be localized to the ER membrane. Here it may be able to sense the activation of ER stress pathways and activate the cytosolic CDK5-JNK signaling cascade, among other possibilities. To test this, we performed immunofluorescence staining for Superdeath in S2 cells and looked for co-localization with known subcellular markers: Calnexin 99A (ER), Golgin-84 (Golgi), and Lamp1 (lysosome). We induced expression of a transgenic Superdeath tagged with the V5 epitope in S2 cells and stained for V5 and each subcellular marker. We found that while Superdeath appears to co-localize with Calnexin 99A (Fig 6A), it does not co-localize with Golgin-84 (Fig 6B). While the vast majority of the Superdeath and Lamp1 signals are distinct and non-overlapping, we do detect a small minority of overlapping signals (Fig 6C). Importantly, in all cases, Superdeath does not appear to localize to the plasma membrane, the primary location of several potential human orthologues such as ANPEP, ENPEP, and LVRN. The staining for V5 is not detectable in cells not expressing the *superdeath* transgene (Fig S2A,B). We concluded that Superdeath primarily localizes and functions at the ER membrane.

**Figure 6.**
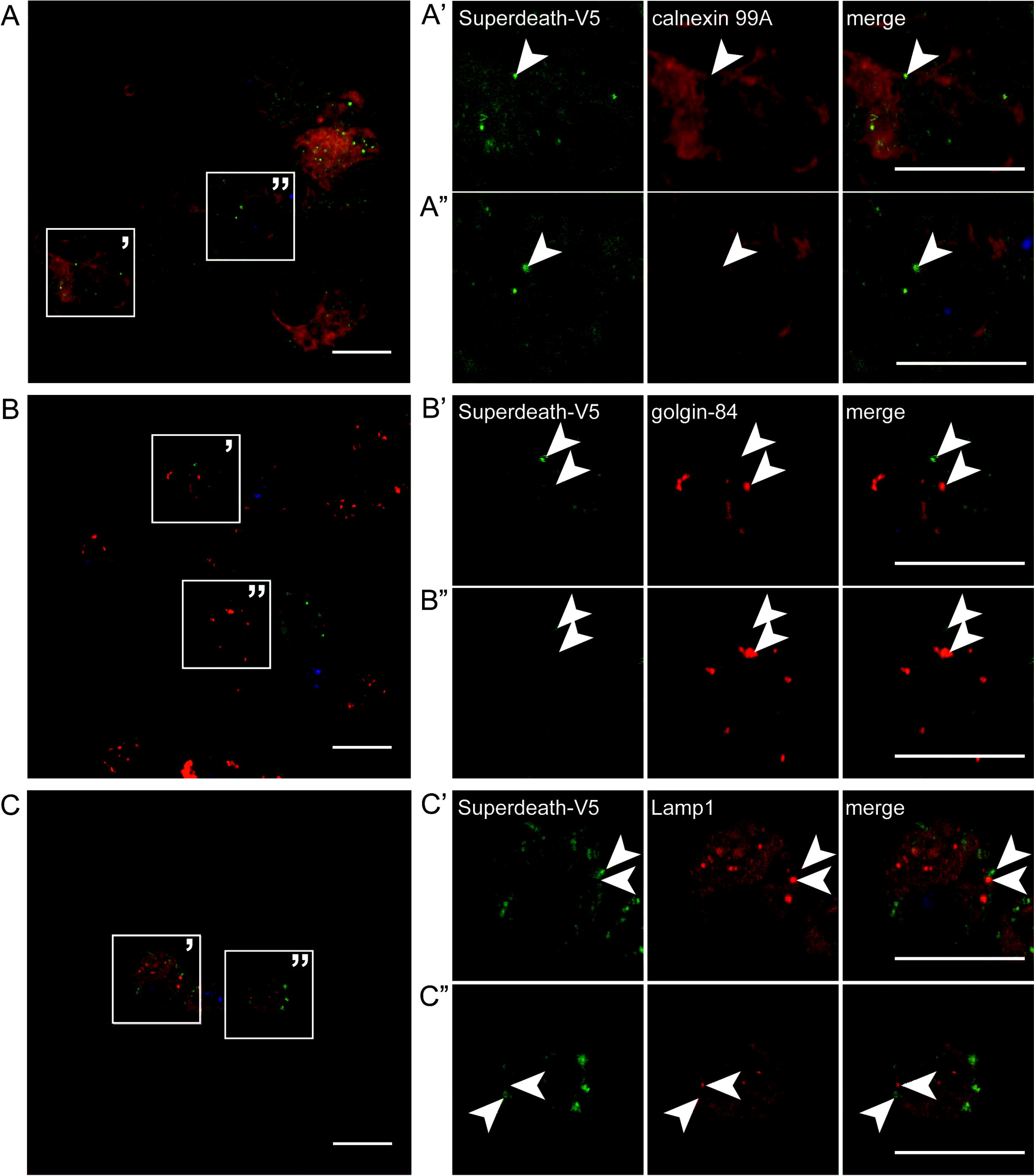
Superdeath is localized to the endoplasmic reticulum. Superdeath predominantly localizes to the ER membrane. **A.** Superdeath localizes to the ER. S2 cells expressing Superdeath-V5 were stained for V5 (green) and Calnexin 99A (red) and counterstained with DAPI. **A’** and **A”** represent the highlighted panels from **A**. White arrows highlight select sites of V5 and Calnexin 99A overlap. **B.** Superdeath does not localize to the golgi. S2 cells expressing Superdeath-V5 were stained for V5 (green) and Golgin-84 (red) and counterstained with DAPI. **B’** and **B”** represent the highlighted panels from **B**. White arrows indicate select sites of independent V5 staining or Golgin-84 staining. **C.** Superdeath does not primarily localize to the lysosome. S2 cells expressing Superdeath-V5 were stained for V5 (green) and Lamp1 (red) and counterstained with DAPI. **C’** and **C”** represent the highlighted panels from **C**. White arrows indicate select sites of independent V5 staining or Lamp1 staining. Scale bars = 0.01 mm.

## DISCUSSION

Activation of the ER stress response and the subsequent cell death is a major contributor to the pathogenesis of a number of human diseases. Degenerative diseases are commonly caused or complicated by the accumulation of misfolded proteins in the ER and stress-induced cell death (2, 32–35). In order to treat these degenerative diseases, it is essential to specifically target stress-associated cell death without inhibiting the beneficial stress-induced pathways that restore homeostasis.

In this study, we examined the metallopeptidase *superdeath*, a modifier of Rh1^G69D^-induced degeneration (9). Loss of *superdeath* activity reduces apoptosis and degeneration without impacting the activation of the ER stress sensors IRE1 and PERK. Both of these sensors, while capable of initiating apoptosis upon chronic activation, have important cell survival functions that are essential for returning the stressed cell to homeostasis (5). Loss of *superdeath* leaves these beneficial functions largely intact, and instead, reduces the activation of JNK signaling through CDK5 activation. In this manner, *superdeath* orthologues could serve as important therapeutic targets to tip the balance in favor of cell survival in degenerative diseases.

Additionally, *superdeath* fills an important gap in ER stress-associated cell death biology. While previous studies have shown that CDK5 is responsible for activating the JNK signaling cascade under ER stress (6), the mechanism of how the stress signal is communicated to the kinase is still unknown. Our data suggest that the ER-associated Superdeath, which is likely orthologous to ERAP1 or ERAP2 in humans, may serve as an important bridge between ER stress and CDK5. Importantly, the active site of Superdeath is predicted to be present on the cytoplasmic side of the ER, suggesting that changes in ER membrane conformation and ER luminal environment could alter Superdeath activity, directly or indirectly activating CDK5.

There is already evidence that ERAP2 could be playing an important role in autoimmune disorders such as Crohn’s disease and inflammatory arthritis, as well as in the response to viral infection (36–38). The types of stress responses induced in these diseases, including ER, oxidative, and mechanical stresses, frequently activate cell death through the JNK signaling cascade. This raises the possibility that Superdeath and ERAP2 are regulating apoptosis in response to many cellular stresses and that these roles are not limited to the induction of ER stress, and this is an exciting avenue of future study.

Our findings suggest that the *Drosophila* gene *superdeath* regulates stress-induced apoptosis. We have demonstrated a role for this gene in known apoptotic pathways and have shown that its role lies downstream of the beneficial activation of the UPR. The position of *superdeath* in the cellular stress response makes its orthologues attractive candidates for therapeutic targeting for a variety of diseases associated with ER stress-induced degeneration. Understanding how modifiers of stress-induced apoptosis are functioning in the cell also increases our understanding of degenerative diseases and provides new avenues for personalized therapies.

## METHODS

### Fly stocks and maintenance

Flies were raised at room temperature on standard diet based on the Bloomington Stock Center standard medium with malt. The strain containing *GMR*-*GAL4* and *UAS*-*Rh1*^*G69D*^ on the second chromosome (*GMR*>*Rh1*^*G69D*^) has been previously described (9, 16). The following strains are from the Bloomington Stock Center: *MS1096*-*GAL4* (8696), UAS-*superdeath* RNAi (42947), control *attP40* (36304). The *UAS*-*CDK5* RNAi is from the Vienna *Drosophila* Resource Center (104491). The puc-LacZ enhancer trap is available from Kyoto (109029). The strains containing the *UAS*-*Xbp1*-*EGFP* transgenes were a gift from Don Ryoo (NYU).

### Eye/wing imaging

For eye and wing images, adult females were collected under CO_2_ anesthesia and aged to 2-7 days, then flash frozen on dry ice. Eyes were imaged at 3X magnification using a Leica EC3 camera. Wings were dissected away from the body, then imaged at 4.5X magnification using the same camera. Eye and wing area were measured in ImageJ as previously described (9).

### Immunohistochemistry

Eye discs were dissected from wandering L3 larvae in cold 1X PBS, then immediately transferred to cold 4% PFA on ice. S2 cells were treated while adhered to sterile plastic coverslips. Samples were fixed in 4% PFA for 15-20 min, then washed in 1XPAT (0.1% TritonX100) prior to blocking with 5% normal donkey serum. Samples were stained with primary antibodies for rhodopsin (1:50, Developmental Studies Hybridoma Bank #4C5), GFP (1:2000, Thermo-Fisher #A6455), LacZ (1:20-1:50, DSHB #40-1a), Calnexin 99A (1:50, DSHB #cnx99A 6-2-1), Golgin-84 (1:50, DSHB #golgin-84 12-1), Lamp1 (1:100, Abcam # ab30687), and V5 (1:500, Cell Signaling #13202S, and 1:250, Thermo-Fisher #R960-25). Apoptosis was monitored using the ApopTag Red *In Situ* Apoptosis Detection Kit (Millipore #S7165). Samples were mounted in Slowfade™ Diamond Antifade Mountant (Thermo-Fisher #S36967) and imaged with an Olympus FV1000 confocal microscope or a Nikon A1 confocal microscope.

### Western blots

Protein was isolated from 10 brain-imaginal disc complexes were dissected from wandering L3 larvae or S2 cells and homogenized in 1X Laemmli/RIPA buffer containing 1X protease inhibitors (Roche cOmplete Mini EDTA-free protease inhibitor tablets) as well as the phosphatase inhibitors Calyculin A and okadaic acid. Equivalent amounts of protein were resolved by SDS-PAGE (10% acrylamide) and transferred to PVDF membrane by semi-dry transfer. Membranes were then treated with either 5% BSA or 5% milk protein block in 1XTBST prior to immunoblotting. Blots were probed with antibodies for P-eif2α (1:1000, abcam #32157), Pan-eif2α (1:500, abcam #26197), and tubulin (1:2000, Developmental Studies Hybridoma Bank #12G10). Blots shown are representative of at least three biological replicates.

### Tunicamycin treatment

Crosses to generate the indicated genotypes were set up on egg caps containing yeast paste. L2 larvae were then treated with either 10 μg/mL Tunicamycin (diluted 1:1000 from a 10 mg/mL stock solution) or 1:1000 DMSO in Schneider’s media for five hours at room temperature. The larvae were then washed in 1XPBS twice and placed on standard media. Viability was determined by survival to pupation. Survival for each genotype was normalized to the DMSO-treated control condition. Each replicate represents 112-130 larvae per genotype.

### S2 cells

DsRNA was generated using the MEGAscript™ T7 Transcription kit (ThermoFisher #AM1334), with primers for *EGFP* (F: TTAATACGACTCACTATAGGGAGACCACAAGTTCAGCGTGTCC and R: TTAATACGACTCACTATAGGGAGAGGGGTGTTCTGCTGGTAGTG) and *superdeath* (F: TTAATACGACTCACTATAGGGAGATCCGGTGGTTAAGGTGTCAAGG and R: TTAATACGACTCACTATAGGGAGAGCCGGAGTTGACGAACATGG). S2 cells were treated with DsRNA against *EGFP* (as a control) or against *superdeath* at a density of approximately 2 × 10^6^ cells/mL in a 24-well plate. Cells were incubated with DsRNA for 4-7 days before being split and treated with either 2mM DTT or DMSO as a control. Cells were treated for four hours for *Xbp1* splicing and P-eif2a measurements and for 7.5 hours for *rpr* measurements. RNA was isolated from cells using the Direct-zol™ RNA Miniprep Kit and used to generate cDNA (Protoscript II, NEB). Protein was isolated from cells as described above.

Knockdown of *superdeath* was confirmed using qPCR (primers: F: ATTCGCAGCAGTTTCCACCAC and R: TTCGTGGCGAACTTGAACAGC). *Xbp1* splicing was evaluated from the cDNA using PCR (primers for *Xbp1:* F: TCAGCCAATCCAACGCCAG and R: TGTTGTATACCCTGCGGCAG). The spliced and unspliced bands were separated on a 12% acrylamide gel and the proportion of these bands quantified using ImageJ software. *rpr* levels were analyzed by qPCR (F: TTGCGGGAGTCACAGTGGAG and R: AATCCTCATTGCGATGGCTTGC). Transcript levels were normalized to *rpl19* (F: AGGTCGGACTGCTTAGTGACC and R: CGCAAGCTTATCAAGGATGG) and compared between matched DMSO or DTT-treated S2 cells.

### Cloning

Superdeath was overexpressed in S2 cells using the TOPO® TA Cloning® Kit (Thermo-Fisher). The coding sequence for *superdeath* was expressed from the pMT-DEST48 inducible expression vector with a C-terminal V5 tag. S2 cells adhered to sterile glass cover slips were made competent using the Calcium Phosphate Transfection Kit (Thermo-Fisher) and expression of the construct was induced with 500 μM CuSO_4_ for 66 hours. Cells were then stained for the V5 tag and other subcellular markers to determine Superdeath protein localization as described above.

### Statistics

Statistics were calculated using R software. P-values were determined using ANOVA for eye size, fluorescence levels, and transcript levels in qPCR. A pairwise T-test was performed for larval tunicamycin treatment. A cutoff of P = 0.05 was used for significance.

## ACKNOWLEDGEMENTS

We thank Drs. Julie Hollien, Aylin Rodan, Anthea Letsou, and Carl Thummel for use of equipment and sharing reagents. We thank Dr. D. Allan Drummond for his tremendous insight on Twitter and the name *superdeath*. This research was supported by an NIH/NIGMS R35 award (1R356M124780) and a Glenn Award from the Glenn Foundation for Medical Research to CYC. CYC is the Mario R. Capecchi Endowed Chair in Genetics. R.A.S.P. was supported by a NIDDK T32 fellowship (5T32DK110966).

---

**Supplementary Figure 1.**
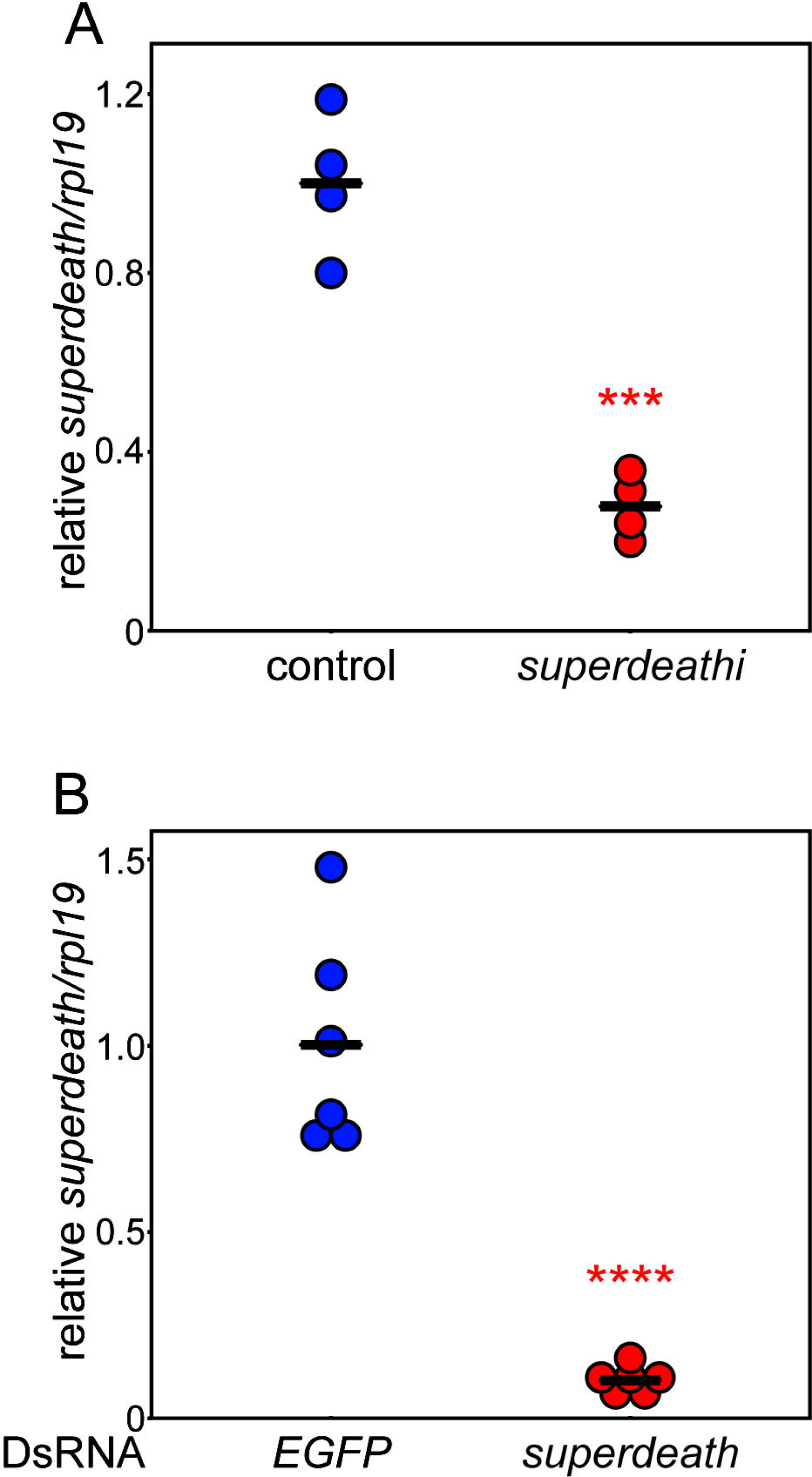
*superdeath* expression is efficiently reduced by both RNAi and DsRNA. The RNAi and DsRNA targeting *superdeath* in our models of ER stress significantly reduce *superdeath* expression. **A.** The Bloomington *Drosophila* Stock Center *superdeath* RNAi strain (42947) used in this study efficiently reduces expression of *superdeath* (27.8% of controls, N = 4). The RNAi construct was driven ubiquitously by *Tubulin*-*GAL4*, and expression determined in wandering L3 larvae compared to controls expressing only *Tubulin*-*GAL4* (N = 4). **B.** S2 Cells treated with DsRNA targeted against *superdeath* also had near complete reduction in *superdeath* transcript levels (10.2%, N = 6) as compared to control cells treated with DsRNA against *EGFP* (N = 6). Values are average ± SD. *** P < 0.0005, **** P < 0.00005.

**Supplementary Figure 2.**
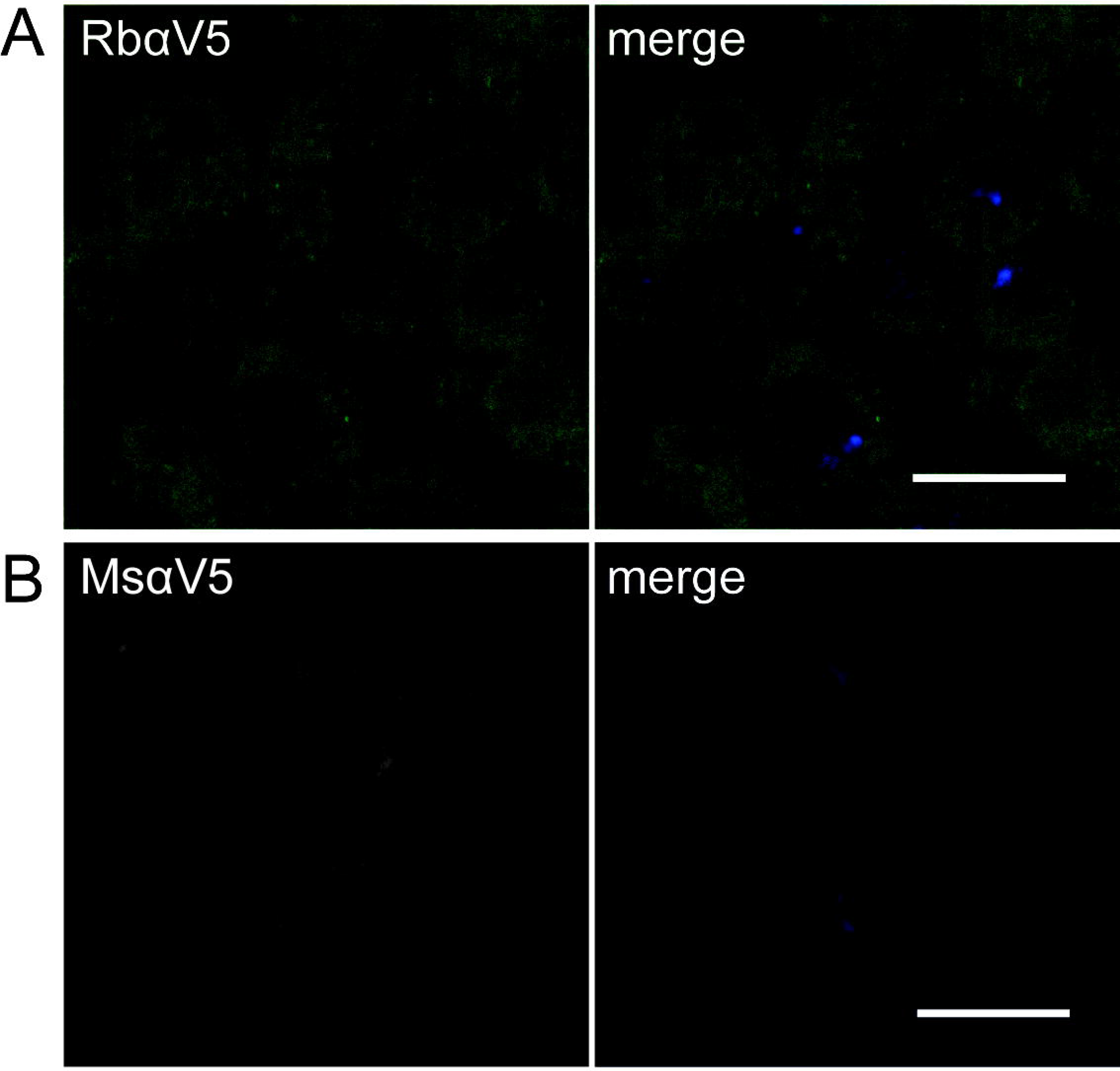
Specificity of V5 staining. The signal for Superdeath-V5 in Figure 6 is specific to expression of the V5 tag. **A.** S2 cells that were not transformed with the pMT-DEST48-Superdeath-V5 vector were stained for rabbit αV5 (green) and counterstained with DAPI. Only background staining and no punctate staining indicative of Superdeath is detectable. **B.** S2 cells that were not transformed with the pMT-DEST48-Superdeath-V5 vector were stained for mouse αV5 (green) and counterstained with DAPI. No punctate staining indicative of Superdeath is detectable. Scale bars = 0.01 mm.

**Supplementary Table 1.** Tunicamycin Sensitivity.

|  | Proportion Survival |  |  |  |  |  |
| --- | --- | --- | --- | --- | --- | --- |
|  | *Tub-GAL4* control |  |  | *Tub-GAL4>superdeathi* |  |  |
|  | DMSO | TUN | TUN/DMSO | DMSO | TUN | TUN/DMSO |
| **Replicate** |  |  |  |  |  |  |
| **1** | 0.900 | 0.800 | **0.889** | 0.913 | 0.846 | **0.927** |
| **2** | 0.833 | 0.742 | **0.890** | 1.000 | 1.000 | **1.000** |
| **3** | 0.952 | 0.867 | **0.910** | 0.933 | 1.000 | **1.071** |
| **4** | 0.929 | 0.794 | **0.855** | 0.917 | 0.917 | **1.000** |
| **Total Treated Larvae** | 113 | 130 |  | 112 | 127 |  |
| **Total Pupae** | 102 | 104 |  | 105 | 119 |  |
| **Proportion Survival** | 0.903 | 0.800 | **0.886** | 0.938 | 0.937 | **0.999** |

